# Sex differences in PRDM9-independent fine-scale recombination patterns

**DOI:** 10.64898/2026.06.19.733336

**Authors:** Julien Joseph

## Abstract

Meiotic recombination rates exhibit strong, fine-scale variation along the genome of many eukaryotes. In some animals, including humans, the protein PRDM9 directs recombination toward so-called recombination hotspots. However, in birds and dogs, which have lost PRDM9, hotspots generally occur in promoter sequences called CpG islands. Furthermore, in many species, recombination rates and patterns differ between the sexes, a phenomenon known as heterochiasmy. While sex differences in recombination rates and broad-scale genomic variation are rather well documented, far less is known about differences in fine-scale variation, with no insight from PRDM9-lacking animals. In this study, using thousands of high-resolution crossovers, I demonstrate that fine-scale recombination patterns are globally similar between the sexes in both dogs and barn owls. However, I show that recombination in protein-coding gene promoters is heavily male-biased in both species. Overall, the shared patterns of fine-scale heterochiasmy between these two amniotes shed light on both the sex-specific regulation of recombination and the evolutionary forces that shape it.

## Introduction

Meiotic recombination is a key evolutionary process shaping the evolution of eukaryotic genomes. During the production of sex cells, meiotic recombination is initiated by the induction of a double-stranded break (DSB) on one of the parental chromosome, which is then repaired by using the other chromosome as a template. This repair either induces an exchange of chromosome arm called Crossover (CO) or only a local transfer of DNA called a Non-Crossover (NCO). This exchange of DNA between parental chromosome generates novel combination of genetic variants, enhancing the genetic and functional diversity on which natural selection can act (Felsenstein, 1974). Furthermore, in most species, meiotic recombination is an essential step for correct chromosome segregation, the success of meiosis and therefore for viable offspring production (Kaback et al., 1992; Hassold et al., 2007).

Within an individual, the CO rate generally varies between and along chromosomes (reviewed in Stapley et al. (2017); Zelkowski et al. (2019); Peñalba and Wolf (2020)). In most species, CO rate is typically higher in short chromosomes and subtelomeric regions and lower in large chromosome and centromeric regions (Haenel et al., 2018; Zelkowski et al., 2019; Brazier and Glémin, 2022). Moreover, in many eukaryotes, CO events are concentrated into hotspots of a few kilobases (Lichten and Goldman, 1995; Petes, 2001; Myers et al., 2005; Tock and Henderson, 2018). In many animals, including humans, these hotspots are directed by the protein PRDM9 that binds specific DNA motifs determined by a zinc finger array (Baudat et al., 2010; Parvanov et al., 2010; Myers et al., 2010; Hoge et al., 2024; Raynaud et al., 2025). PRDM9 arose in the ancestors of all animals (Oliver et al., 2009; Ponting, 2011), but has been lost in many lineages, including birds and canids (dogs, wolves and foxes) (Axelsson et al., 2012; Singhal et al., 2015; Baker et al., 2017; Cavassim et al., 2022). In species which have lost PRDM9, hotspots are rather concentrated in short sequences called CpG islands (CGIs) (Auton et al., 2013; Singhal et al., 2015; Kawakami et al., 2017; Topaloudis et al., 2025). CGIs are regulatory sequences characterized by low levels of DNA methylation (Bird, 1980). This low methylation level prevents spontaneous deamination of methyl-CpG dinucleotides into TpGs, leading to a high density of CpGs compared to the rest of the genome. Moreover, in vertebrates, CGIs almost always colocalize with H3K4me3 histone marks, a universal mark of transcription initiation (Hughes et al., 2020). Indeed, in humans and mice, a large fraction of CGIs act as promoter for protein coding genes, mostly housekeeping genes (Deaton and Bird, 2011). On the other hand, many CGIs are located far from the promoter of protein-coding genes, and are called orphan CGIs (Illingworth et al. (2010); Fig.1). These orphan CGIs are often involved in regulatory functions by initiating the transcription of non-coding RNAs, but the functional relevance of many of them remains unclear (Illingworth et al., 2010; Deaton and Bird, 2011). Moreover, to this day, the exact mechanism by which CGIs are linked to the DSB machinery in meiosis remains unknown (Joseph et al., 2026). One possibility is that DSBs are specified by H3K4Me3 histone marks, like in yeasts (Sommermeyer et al., 2013; Acquaviva et al., 2013; Adam et al., 2018). Another possibility is that they are specified directly by DNA methylation through recognition of non-methylated cytosines (Joseph et al., 2026).

**Figure 1:**
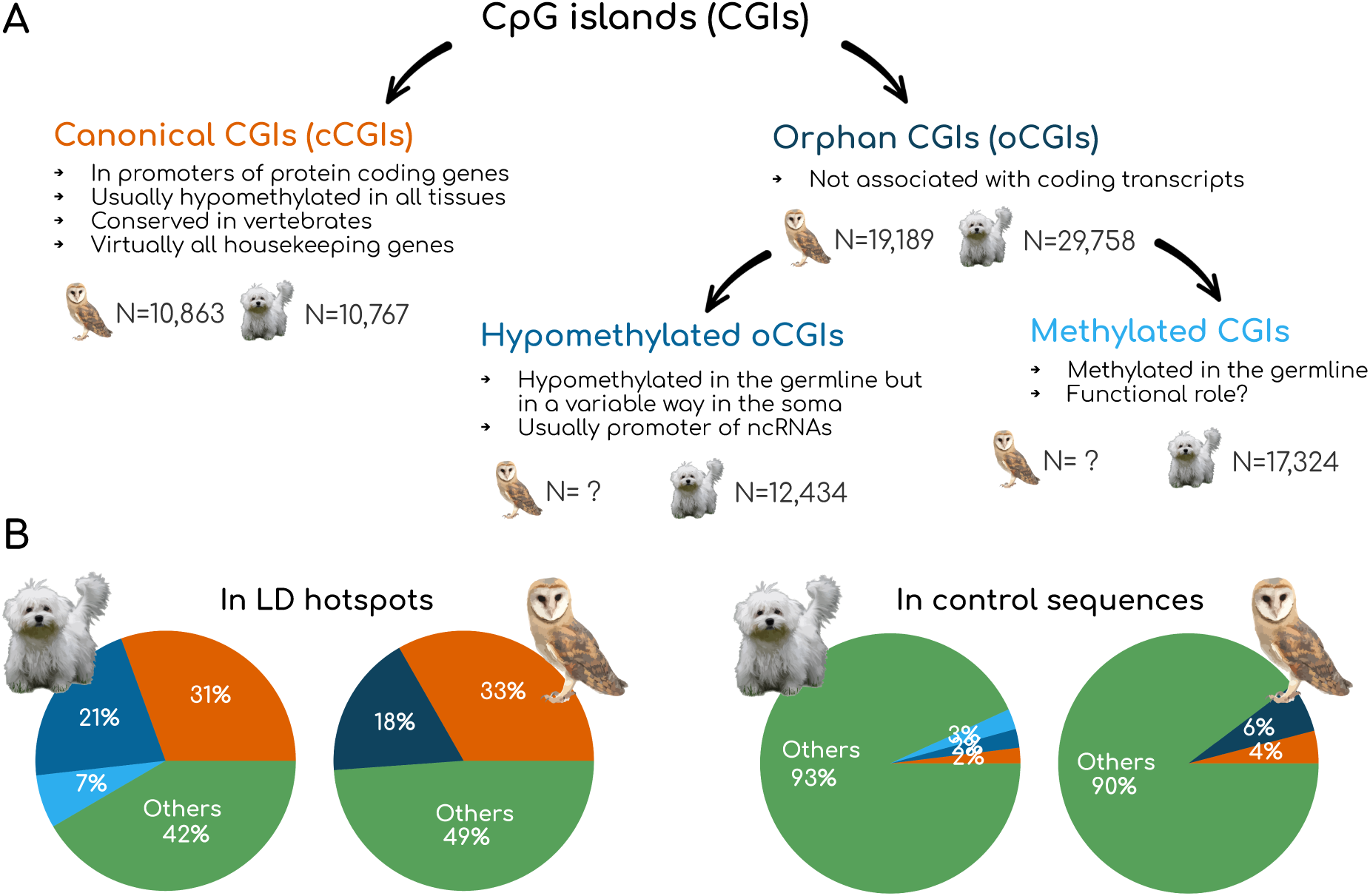
The different types of CGIs and their overlap with LD-based hotspots. A) Classification of CGIs used in this study with some key characteristics summarized in Deaton and Bird (2011). B) Overlap between LD-based hotspots and the different categories of CGIs presented in A) in dogs and barn owls. Control sequences were obtained by randomly shuffling hotspots, including only regions in which CGIs can be called (see Material and Methods).

In addition to variation within the genome, the CO rates and patterns of many species vary between the sexes, a phenomenon known as heterochiasmy. Typically, CO rate is higher and more homogeneous in females while in males, recombination rate is often lower and concentrated in subtelomeric regions, but many exceptions exist (reviewed in Sardell and Kirkpatrick (2020)). These observations have fueled several theories explaining these sex differences from molecular and evolutionary perspectives (Burt et al., 1991; Lenormand, 2003; Lenormand and Dutheil, 2005; Mank, 2009; Haig, 2010; Brandvain and Coop, 2012; Blackmon and Brandvain, 2017; Sardell and Kirkpatrick, 2020). Nevertheless, none of these theories fully explains the observed patterns of heterochiasmy across eukaryotes. Sex differences in recombination rates and patterns therefore remain one of the most enigmatic conundrums in evolutionary biology (Lenormand et al., 2016; Sardell and Kirkpatrick, 2020). Interestingly, the most recent evolutionary theories make important predictions regarding sex-differences in fine-scale recombination patterns, particularly around gene promoters (Sardell and Kirkpatrick, 2020). However, with the exception of a few model species, our knowledge of sex differences in fine-scale CO patterns remains limited. Indeed, mapping a large number of sex-specific COs at high resolution requires sequencing the entire genome of hundreds of related individuals, a process that remains very costly.

In humans, both sexes appear to share similar usage of CO hotspots directed by PRDM9 (Coop et al., 2008; Bhérer et al., 2017). However, women exhibit an elevated CO rate around promoters (Bhérer et al., 2017). Furthermore, the CO rate differs between the sexes in certain families of repeated elements (Bhérer et al., 2017). In mice, the usage of double-stranded break (DSB) hotspots vary between the sexes, but it appears that this variation is mainly due to sex differences in regional DSB rate, with a more complex contribution of DNA methylation patterns (Brick et al., 2018). Outside of animals, in maize, Kianian et al. (2018) showed that both sexes recombine strongly in promoters and close to transcription ending sites, but no sex difference in magnitude has been reported. On the other hand, nothing is known about fine-scale heterochiasmy in animals that have lost PRDM9.

Here, using whole genome resequencing data of large pedigrees, I mapped ∼5,000 high resolution COs in the dog genome. I then compared the fine-scale, sex-specific distribution of these COs to that of ∼7,500 published CO events mapped in the barn owl genome, which also lacks PRDM9. I show that both sexes use similarly recombination hotspots defined by fine-scale recombination maps based on linkage disequilibrium (LD). However, both species show a strong male bias in the usage of CGIs associated with the promoters of protein-coding genes. I show that this male bias cannot be explained only by global differences in DNA methylation between promoter-associated and other CGIs across the male germline. Finally, despite recombination enrichment in certain families of repeated elements, no significant sex differences in recombination rates could be detected.

## Results

### Fine-scale mapping of COs in dogs

In vertebrates, several studies demonstrated that recombination rate is higher in promoters and more specifically CpG islands (CGIs) (Auton et al., 2013; Singhal et al., 2015; Kawakami et al., 2017; Topaloudis et al., 2025). However, whether this pattern is driven by recombination in one sex is currently unknown. A previous study attempted to address this question by estimating sex differences in recombination rates in promoters and CGIs in dogs, but did not recover any local increase in females, and only a 15% increase in males (Figure S15 in Campbell et al. (2016)). Unfortunately, the sex-specific crossovers (COs) used in Campbell et al. (2016) were obtained by genotyping a pedigree of 237 dogs at 163,400 SNPs, and lacked sufficient resolution to tackle this question (see fig. S2). Therefore, I used the whole genome sequencing data from a pedigree of 404 dogs (mean coverage of 40X) from Zhang et al. (2025), to map CO events at a fine scale. I was able to map 17,036 sex-specific COs in the dog genome using the approach of Coop et al. (2008) (details in Material and Methods). To confirm the accuracy of the CO inference, I compared the broad-scale CO distribution with that of Campbell et al. (2016). In 5Mb windows, sex-averaged recombination rates estimated in this study (JJ) are highly correlated to those of Campbell et al. (2016) (CC) (Pearson R=0.71, fig. S1A).

Moreover, sex-specific recombination rates correlate better between studies than between sexes (fig. S1B), indicating that the two methods obtain coherent results, even on different datasets. Unfortunately, because parents from the dataset of Zhang et al. (2025) are from the same dog breed, many of the COs called in this study occur in regions of high homozygocity, and hitherto low marker density (see Material and Methods). Nevertheless, with this new dataset, enough CO events have sufficient resolution to address the question of their fine-scale distribution (fig. S2). I therefore, filtered this dataset to obtain COs whose inter-marker distance is inferior to 10 kb, which represents 5,013 high resolution COs (see fig. S1).

### Sex-specific CO hotspot usage in dogs and barn owls

Recombination hotspots have previously been identified using sex-averaged LD-based recombination maps in both dogs and barn owls (Auton et al., 2013; Topaloudis et al., 2025). However, it is possible that only one sex recombines in hotspots, with the other not using them. Alternatively, these hotspots could be shared between the sexes or the LD hotspots could comprise male- and female-specific/biased hotspots. To test these hypotheses, I first examined the elevation of the CO rate in LD hotspots compared to their flanking sequences in both dogs and barn owls. Comparing CO rate to flanking sequences ensures that the recombination elevation do not capture regional variation in recombination rates. This is necessary to remove sex-biased recombination in hotspots that can be due to broad-scale heterochiasmy. In both species, there was a similar CO increase in LD hotspots for both sexes (Fig.2). However, if hotspots occurring in CGIs associated with the promoter of protein-coding genes (cCGIs) are distinguished from those occurring in orphan CGIs (oCGIs) and from those occurring outside CGIs, sex differences start to be noticeable (Fig.2B&D). In both species, recombination in cCGIs is strongly male biased, with almost a fourfold sex difference in dogs and a twofold sex difference in barn owls (Figure.2B&D).

**Figure 2:**
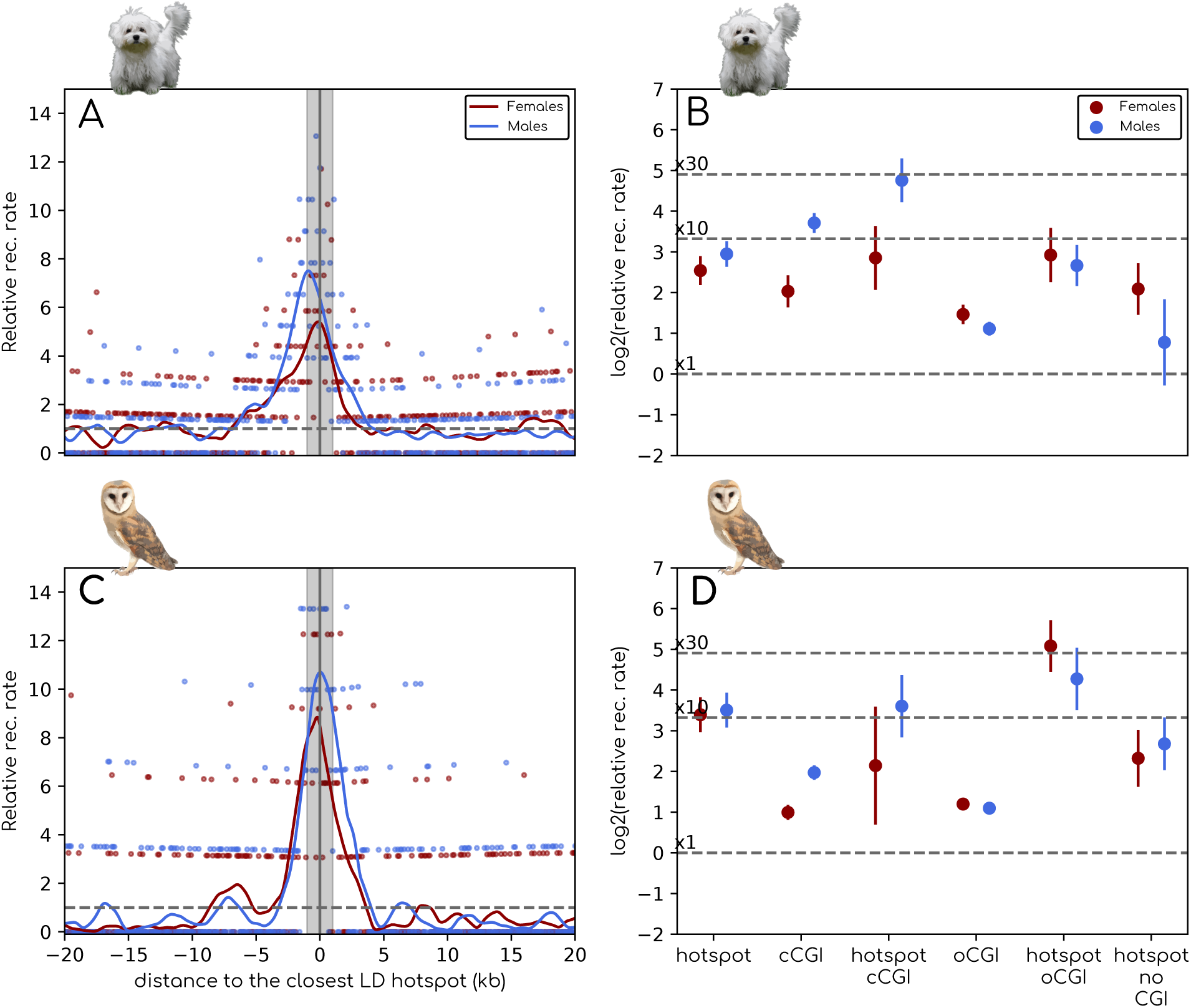
A&C) Relative recombination rate as a function of the distance to the center of the closest LD hotspot in dogs (A) and in the barn owl (C). Relative recombination rate is computed as the number of COs per base pair in a 100bp window divided by the number of COs per base pair in the regions from 5 to 30kb upstream and downstream of the hotspot center. Each point corresponds to a 100bp window, and the lines show a smoothing of the data using a loess function. Blue lines and points correspond to males and red ones to females. B&D) log_2_ of the relative recombination rate in features versus flanking regions in dogs (B) and barn owls (D). Relative CO rate was computed as the ratio between the number of COs per base pairs in a 3kb region around the feature center and the number of COs per base pairs in the flanking regions defined as above. Error bars correspond to 95% confidence interval obtained by bootstrapping CO events (see more details in Material and Methods) On the other hand, no significant sex difference could be noticed in oCGIs or in non-CGI hotspots in both species (Fig. 2B&D).

### DNA methylation and male-biased hotspots in dogs

CGIs are determined by sequence characteristics that are hallmarks of the absence of DNA methylation: a higher density of CpG dinucleotides compared to the rest of the genome. In fact, several studies have found strong associations between recombination rate and methylation levels across eukaryotes (Yelina et al., 2015; Wallberg et al., 2015; He et al., 2017; Joseph et al., 2024). However, high CpG density may reflect past methylation patterns and it is not guaranteed that CpG rich sequences are currently hypomethylated and carry associated H3K4Me3 marks. In fact, many of sequence-based CGIs are found to be methylated in sperm (Cohen et al., 2011; Qu et al., 2018). The functional role and evolutionary origin of these methylated CGIs is unclear (Cohen et al., 2011; Berglund et al., 2015).

Importantly, these sperm methylated CGIs are found almost exclusively outside of promoter regions (in oCGIs). It is therefore possible that the differences in male CO rate observed between cCGIs and oCGIs reflect differences in male germline DNA methylation/H3K4Me3 levels. To test this hypothesis, using hypomethylated regions (HMRs) called in dog sperm in (Qu et al., 2018), I distinguished methylated oCGIs from hypomethylated ones. CGIs that are methylated in sperm indeed show a lower recombination rate compared to hypomethylated ones in both sexes, but still show a twofold increase in female (Fig. 3C).

**Figure 3:**
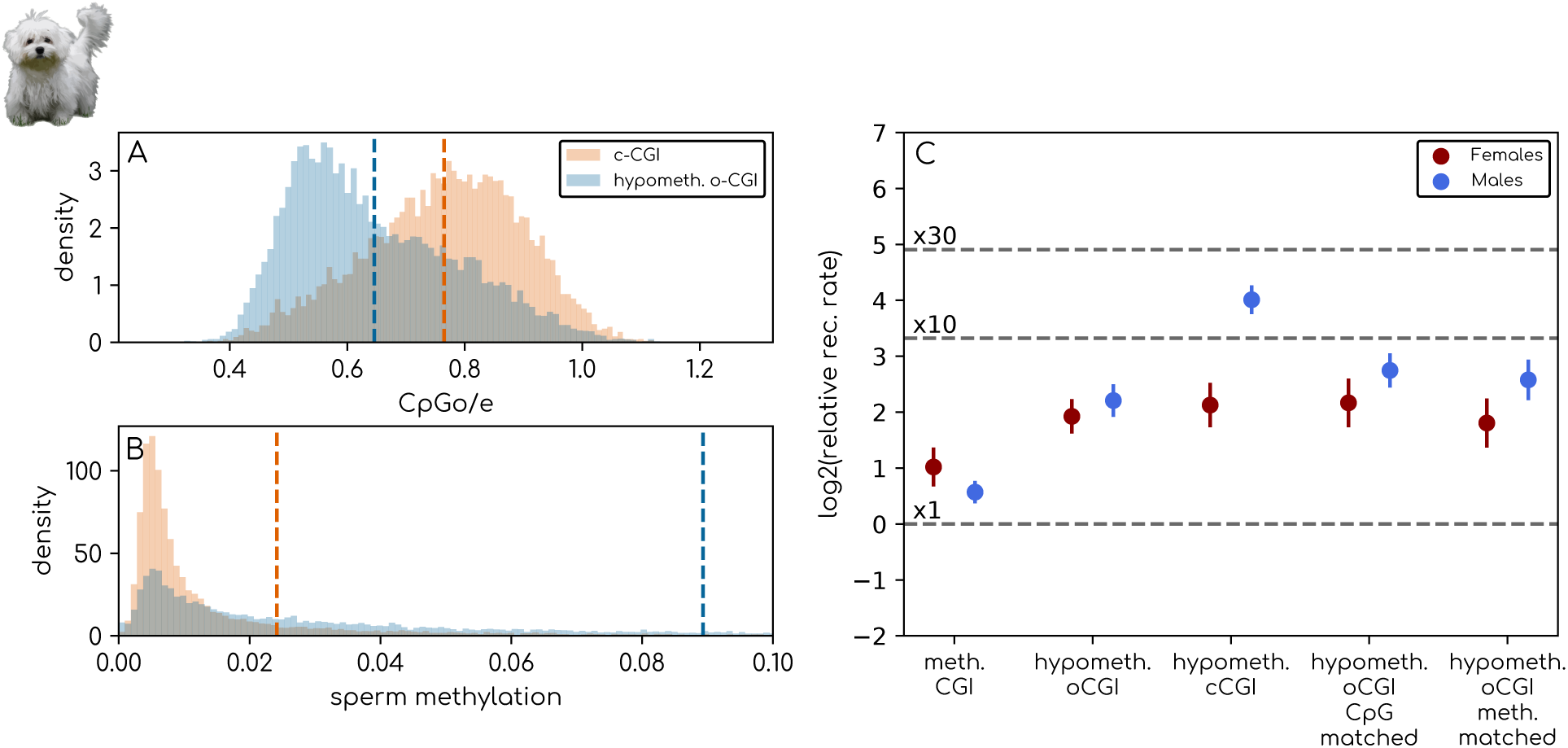
A) Distribution of CpG*_o/e_* ratio in the dog’s hypomethylated oCGIs and cCGIs. CpG*_o/e_* is computed as the number of CpG dinucleotides divided by the product between the number of guanine and cytosine in a 3kb region centered around the CGI. B) distribution of dog sperm methylation level in hypomethylated oCGIs and cCGIs. Methylation level is the percentage of methylated CpG dinucleotides in in a 3kb region centered around the CGI. Percentage of methylation of individual CpG dinucleotides were obtained from the bisulfite sequencing experiment of Qu et al. (2018). C) log_2_ of the relative recombination rate in features versus flanking regions in dogs. Relative CO rate was computed as the ratio between the number of COs per base pairs in a 3kb region around the feature center and the number of COs per base pairs in the flanking regions defined as above. Error bars correspond to 95% confidence interval obtained by bootstrapping CO events (see more details in Material and Methods). CpG matched CGIs are a set of oCGIs selected so that their CpG*_o/e_* distribution matches that of cCGIs. Meth. matched CGIs are a set of oCGIs selected so that their sperm methylation level distribution matches that of cCGIs.

When excluding these sperm-methylated CGIs, we can see that oCGIs do not reach the same CO rate as cCGIs (Fig. 3C). This could be explained by the fact that despite being globally hypomethylated, those CGIs still have higher levels of DNA methylation compared to cCGIs (Fig. 3A&B). I therefore selected a set of oCGIs so that the distribution of either CpG*_o/e_* (a proxy of sex-averaged germline DNA methylation) or sperm methylation level match those of cCGIs. CpG*_o/e_* is computed as the number of CpG dinucleotides divided by the product between the number of guanine and cytosine. CpG*_o/e_* is equal to one under a total absence of DNA methylation and decreases with the level of DNA methylation. In this control set, we can see that recombination rate is higher, but still three times lower than that of cCGIs (Fig. 3C).

### On the nature of hotspots found outside CGIs

In both dogs and barn owls, many CO hotspots fall outside CGIs. Until now, their significance was difficult to assess given the high number of false positives of LD-based methods (Raynaud et al., 2023). Here, I confirmed that these LD hotspots are genuine CO hotspots (Fig. 2B&D), but whether they correspond to a new class of hotspots or whether they are missed CpG island/hypomethylated regions remains unclear. Therefore, I compared the CpG*_o/e_* distribution of these hotspots to that of CGI hotspots and random sequences outside of CGIs in both dogs and barn owls. CpG*_o/e_* is clearly lower in non-CGI hotspots than in CGI hotspots, yet slightly higher than in randomly drawn sequences (fig. S3A&B). Therefore, it is possible that this slight enrichment reflects the fact that some of these sequences are actually CGIs, which would explain their high recombination rate. To test that, I matched the CpG*_o/e_* distribution of these sequences to that of the randomly drawn sequences. In this control dataset, we still see a significant enrichment of recombination in female dogs and in both sexes in the barn owls (fig. S3C). This shows that the increase of recombination rate in these sequences cannot be attributed to the effect of arbitrarily thresholding the CpG*_o/e_* ratio used to define CGIs.

However, these sequences may be very young hypomethylated regions that have not accumulated enough CpG dinucleotides to be classified as CGIs based on sequence criteria. This hypothesis is not supported for the male dog germline, as non-CGI hotspots are largely methylated in sperm (fig. S3D). Nevertheless, male dogs are the only ones that do not exhibit significantly increased recombination rate in non-CGI hotspots (Fig. 2B). Therefore, it is possible that these sequences correspond to new CGIs specific to females in dogs and shared between the sexes in barn owls.

### Sex-specific CO rate in repeated elements

Aside from CGIs, repeated elements have been shown to impact the recombination landscape, mainly because their methylated status represses recombination in their vicinity (Kent et al., 2017). However, in human and mice, some families of repeats are sometimes associated with an increase in recombination rates (Myers et al., 2005). This is mainly due to the fact that PRDM9 can target binding motifs that are enriched in some repeat families (Buard et al., 2014; Yamada et al., 2017; Joseph et al., 2026). In contrast, in dogs, LD hotspots generally occur away from repeated elements, except for GC and CpG rich simple and low complexity repeats, characteristic of CGIs (Auton et al., 2013) The same is true for the DSB hotspots of a Prdm9−/− mouse (Joseph et al., 2026).

Interestingly, in humans, the enrichment of recombination in THE1B transposable elements appears to be mostly male-driven (Bhérer et al., 2017). In turn, we currently don’t know how CO rate in repeated elements differs between the sexes in PRDM9-lacking species. I therefore looked at the sex-specific CO rate in the most abundant repeat families in the genome of both dogs and barn owls.

In dogs, no fine-scale depletion of CO events is observed for all of the repeat families, except for L1 repeats (Fig. 4A). This suggests that targeting hypomethylated regions deviates COs away from repeat-rich regions but not specifically from inside repeats. Moreover, there is no sex differences of CO rate in repeats, except for simple repeats and low complexity repeats where there is both a CO enrichment and a male bias (Fig. 4A). Because these repeats are often found in CGIs (Auton et al., 2013), the CO male-bias in these repeats may be due to the bias found at cCGIs. Indeed, after masking CGIs from repeats, the male bias disappears (Fig. 4C).

**Figure 4:**
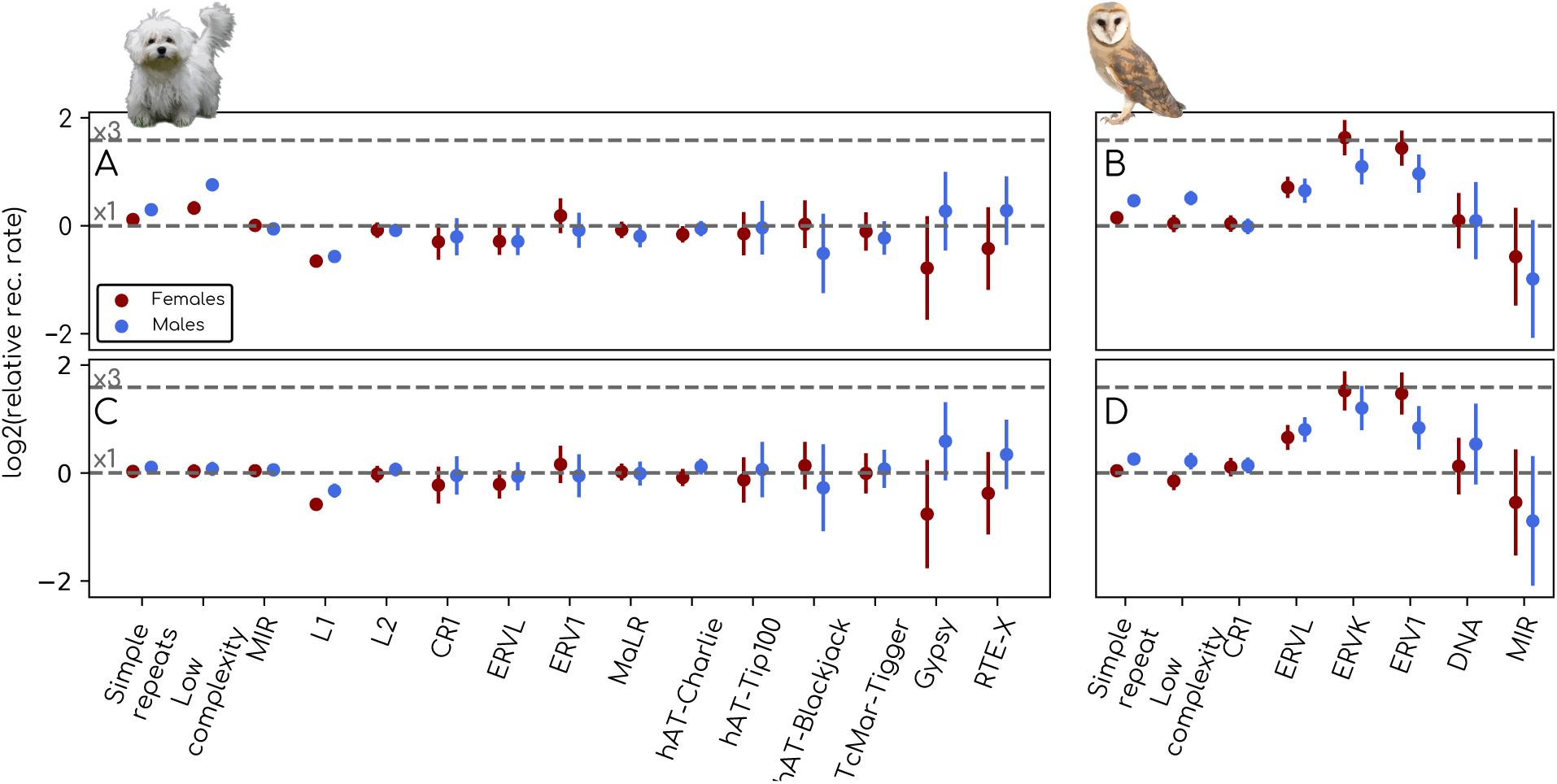
A&B) log_2_ of the relative recombination rate in repeated elements versus flanking regions in dogs (A) and barn owls (B). Relative CO rate was computed as the ratio between the number of COs per base pairs falling in a given repeat family, including the 1.5kb flanking regions, and the number of COs per base pairs in the 5 to 30 kb flanking regions defined as above (see Material and Methods). C&D) Same as A&B after removing the 3kb around the center of CGIs from both repeat families and their flanking regions (see Material and Methods). Error bars correspond to 95% confidence interval obtained by bootstraping CO events (see more details in Material and Methods)

Barn owls do not show fine-scale depletion of CO events in repeats either (Fig. 4B). Surprisingly, however, there is a local increase in CO rates in endogenous retroviruses (ERVs), with an approximately threefold increase in ERVLs and ERVKs in female, higher than that of CGIs (Fig. 4B). More puzzling still, masking CGIs from these repeats does not eliminate the local increase, demonstrating that these elements are recombination hotspots *per se* in barn owls (Fig. 4D). In fact, 52% of hotspots that are outside CGIs are at less than 2kb away from an ERV element, while we expect 12.5% by chance. Finally, except for simple and low-complexity repeats, no significant sex difference in recombination rates was detected (Fig. 4B&D^).

## Discussion

### On the mechanisms of PRDM9-independent hotspot determination

Although recombination events are concentrated into hotspots in many eukaryotes, the differences in hotspot usage and positioning between sexes remain largely unexplored. In this study, I showed that fine-scale recombination patterns are similar between the sexes in two distantly-related amniotes lacking the gene PRDM9.

Unfortunately, the mechanism by which PRDM9-independent DSBs are specified is largely unknown in vertebrates (Joseph et al., 2026). One possibility is that these DSBs are specified by H3K4Me3 histone marks, as they are in yeast (Sommermeyer et al., 2013; Acquaviva et al., 2013; Adam et al., 2018). Another possibility is that they are directly mediated by DNA methylation through the recognition of demethylated CpGs or, conversely, by deviation from methylated cytosines (Joseph et al., 2026). Importantly, the genome of female mammals is almost entirely demethylated during meiotic prophase I, when DSBs are initiated. If DSBs were directly regulated by DNA methylation, then female dogs should recombine everywhere and should largely lack recombination hotspots. However, the fact that female dogs also use CpG islands as hotspots rather suggests that DSBs are directed toward histone marks, probably H3K4Me3, which are present at CpG islands in both males and females (Deaton and Bird, 2011; Hanna et al., 2018). The maintenance of these marks at CpG islands through oocyte demethylation is likely crucial to guiding the re-establishment of DNA methylation in later oogenesis stages.

Interestingly, in both dogs and barn owls, hotspots located in gene promoters are strongly male-biased. In dogs, I investigated whether differences of male germline DNA methylation between canonical and orphan CGIs could account for this male bias. It is thought that methylation patterns in males are broadly conserved within the germline, including sperm cells, although not necessarily in their finer details (Lindsay et al., 2019; Greenberg and Bourc’his, 2019). I demonstrated here that the male-specific increase in recombination rate in coding CGIs cannot be explained by patterns of sperm methylation or CpG density alone. The latter reflects the methylation status of the sequence integrated over the entire germline. This could suggest a stronger demethylation associated with an additional deposition of CGI-associated epigenetic marks (such as H3K4Me3) mark at promoter CGIs specifically during meiotic prophase, increasing the probability of realizing DSBs at these loci. Under this hypothesis, recombination does not fall “by default” in regions which already bear CGI-associated epigenetic marks, but that some of these marks are deposited during meiosis only, similarly to the histone marks deposited by PRDM9 (Brick et al., 2012; Eram et al., 2014; Powers et al., 2016; Davies et al., 2016; Grey et al., 2017). Alternatively, this excess of recombination could be due to DNA methylation independent recruitment of CGI-associated epigenetic marks at promoters in the male germline specifically.

I demonstrated that numerous PRDM9-independent hotspots were located outside of CGIs. These hotspots may correspond to regions that have recently become hypomethylated and have not accumulated enough CpG dinucleotides to be classified as a CGI based on their sequence. In barn owls, these hotspots are frequently located within ERV transposable elements. This could suggest that these elements have successfully evaded mechanisms of TE repression, leading to the accumulation of H3K4me3 marks in these repeats, as observed in mutant mice deficient for germline TE repression mechanisms (Zamudio et al., 2015). ERVs could therefore represent a strong risk of deleterious ectopic recombination in this species (Charlesworth et al., 1997; Langley et al., 1988; Maloisel and Rossignol, 1998; Kent et al., 2017; Joseph et al., 2026).

### Species with PRDM9

It has been noted that PRDM9-independent recombination hotspots can coexist with PRDM9-directed ones in species with PRDM9 (Smagulova et al., 2016; Hoge et al., 2024; Joseph et al., 2024), with some differences between the sexes (Brick et al., 2018). In humans, female recombination rates are higher in promoters, but not in males (Bhérer et al., 2017). However, this increase occurs at a much larger scale (∼40kb around the promoter), despite sufficient resolution to resolve the position of CO events (Bhérer et al., 2017). Moreover when considering only promoter far from PRDM9 binding motifs, this increase disappears (Bhérer et al., 2017). It is therefore likely that this increase is more due to sex-biased usage of PRDM9 hotspots around promoter (at a larger scale), rather than a sex-difference in PRDM9-independent recombination hotspot usage. In mice, females tend to use more PRDM9-independent DSB hotspots (Brick et al., 2018). However, whether this is true in cCGIs but not in oCGIs is unclear and is difficult to test quantitatively due to the low overall usage of these DSB hotspots (Brick et al., 2018). This is further complicated by the fact that PRDM9-independent DSB hotspots in mice have been defined in a male PRDM9 knockout, with no information from females. Overall, sex-differences in recombination hotspot usage in vertebrates have only been investigated in species with very low usage of PRDM9-independent recombination hotspots (Joseph et al., 2024), which complicates their investigation. Future studies in species with higher usage of PRDM9-independent hotspot could shed light on whether the heterochiasmy patterns recovered in this study are also present in species with a functional PRDM9.

### Evolutionary significance

Sex differences in recombination rates and patterns are one of the most puzzling mysteries revolving around the evolution of sex in eukaryotes (Lenormand et al., 2016; Sardell and Kirkpatrick, 2020). On the one hand, it is possible that heterochiasmy only results from the different ways meiosis is performed in spermatocytes versus oocytes (Lenormand et al., 2016; Sardell and Kirkpatrick, 2020; Johnston, 2024). However this hypothesis does not explain why patterns of heterochiasmy can vary between closely related species with very similar gametogenesis (Mank, 2009; Sardell and Kirkpatrick, 2020). Another theory, coined SACE (sexually antagonistic cis-epistasis), proposes that the occasional suppression of recombination between promoters and their cis-regulatory elements can be selected for in the sex in which sexual/haploid selection is strongest (Sardell and Kirkpatrick, 2020). This is explained by the potential for stronger epistasis between promoters and cis-regulators in this sex, which would favour tight genetic linkage (Sardell and Kirkpatrick, 2020). In mammals and birds, this hypothesis predicts a lower recombination rate in male promoters. Here, I showed that on the contrary, recombination at promoters is generally two to four times higher in males versus females in two distantly related-amniotes. This suggests that indirect selection on recombination rates mediated by sexually antagonistic selection is not the main driver of fine-scale heterochiasmy in PRDM9-lacking species. However, it remains possible that SACE plays a role in modulating this male bias in species with different strength of sexual/haploid selection.

Notably, the patterns of broad-scale heterochiasmy differ greatly between dogs and barn owls. In dogs, broad-scale heterochiasmy follows the canonical model of male-biased recombination in subtelomeric regions and female-biased recombination in pericentromeric regions (Campbell et al., 2016). In contrast, broad-scale heterochiasmy in barn owls is very limited (Topaloudis et al., 2025). The conservation of fine-scale heterochiasmy in these two species suggests that fine-scale heterochiasmy in amniotes lacking PRDM9 might be driven by shared gametogenesis-related constraints yet to be discovered. Broad-scale heterochiasmy, however, would be less constrained and more free to evolve (Burt et al., 1991). Further research on broad-and fine-scale heterochiasmy in various species should clarify the generality of this pattern.

## Material and Methods

### Cross-over maps

High resolution sex-specific CO positions in the barn owl were retrieved from Topaloudis et al. (2025). This includes 3,146 male COs and 3,329 female COs. Low resolution sex-specific COs in dogs, obtained by genotyping a pedigree of 237 dogs at 163,400 SNPs were taken from Campbell et al. (2016) This represents 3,584 male COs and 4,265 female COs. Furthermore, I used the whole genome sequencing data from a pedigree of 404 dogs (mean coverage of 40X) from Zhang et al. (2025), to map CO events at a fine scale.

COs were called using the procedure described in Coop et al. (2008). For male COs, informative markers are loci in which the males are heterozygous, and females are homozygous, and the reverse for female COs. For each of the 86 multisibling families, one offspring is used as a reference. For all other offspring, if they have the same genotype as the reference individual, the marker is annotated with a “1”, and a “2” otherwise. At male-informative markers, every switch between a “1” and a “2” is a potential haplotype switch in the father-inherited chromosome, and conversely for female-informative markers. However, many of these switches correspond to genotyping errors. I therefore excluded all positions with a coverage below 10X in any of the individuals (parents and offsprings). I only included markers for which the heterozygous parents had the allele with lowest coverage supported by at least five reads. Conversely, I only included markers for which the homozygous parent had no reads supporting an alternative allele. For a CO to be called, I required the switch to be supported by at least ten consecutive markers. In some cases, the switch was not clear cut, with consecutive oscillations between the two genotypes. In this case, the lower bound of the CO was defined as the first switch, and the upper bound the last switch before stabilization of the new genotype (at least ten consecutive markers). In families with more than two offspring, if the switch occurred in only one individual, the CO was attributed to it. Switch that occurred at the same marker in all the non-reference individual were called as a CO in the reference individual. When families contained only two offsprings, COs were not attributed to any of the offspring. I then filtered out COs that were at less than 200kb from each other within the same individual (or in any individual in two-sibling families), as they are often artifactual (Coop et al., 2008; Prentout et al., 2025). Only autosomes were considered. The implementation of this procedure can be found at https://gitlab.in2p3.fr/julien.joseph/sexhotuse.

### Calling LD hotspots

Recombination hotspots are usually defined as short sequences with a recombination rate significantly higher than their regional background. In dogs, recombination hotspots had already been called with a method called LDhot, which uses maximum likelihood to call hotspots based on the local increase in recombination rate (Wall and Stevison, 2016). In owls, LD hotspots were defined as the 5% of 2kb windows with the highest recombination rates (Topaloudis et al., 2025). This latter procedure does not take into account the background recombination rate, and may miss hotspots in regions with low recombination rates, or falsely infers hotspots in regions of high recombination rates. Unfortunately, LDhot specifically calls hotspots from outputs of the method LDhat, and cannot be used on the recombination rates estimated in barn owls. To ensure homogeneity between the definition of hotspots in the two species, I called hotspots as 1 kb sequences with a recombination rate higher than their 60kb regional background using a sliding window approach. In dogs, 52% of these 5,698 hotspots recovered with this approach overlap the 6,287 hotspots called by LDhot. Importantly, using either category of hotspots in dogs did not affect the pattern found in this study (fig. S4).

On the other hand, using a tenfold threshold resulted in very few hotspots called in the barn owl (489). Indeed, the relative magnitude of LD elevation in hotspots depend on many parameters (e.g. genetic diversity, demographic history, ratio between recombination and mutation rates), and are not easily comparable between studies (Raynaud et al., 2023; Topaloudis et al., 2025). I therefore lowered the threshold in the barn owl to a fivefold increase that led to 1,794 hotspots. As an illustration of the point explained above, the local increase in recombination rate in these LD hotspots based on pedigree-based crossover mapping is higher in owl hotspots than in dogs (Fig. 2A&B).

### Defining CpG islands

CpG islands are regulatory sequences devoid of DNA methylation characterized by a high GC and CpG content. In the absence of methylation data, they are usually defined as regions with a number of CpG dinucleotides observed over expected (CpG*_o/e_*) higher than 0.6 and a GC content higher than 0.5. CpG*_o/e_* is computed as the number of CpG dinucleotides divided by the product between the number of guanines and cytosines. Dog CGIs are available on the USCS genome browser, called with *cpg lh*, a score-based unpublished algorithm. However they are not available for barn owls. I therefore called CpG islands in both genome using a sliding window approach. I computed CpG and GC content in a 1kb window, sliding with a step size of 50bp. For each of these windows, if the CpG*_o/e_* and GC content are higher than the thresholds mentioned above, the 50bp at the center is considered to be part of a CGIs. Then all positive 50 bp windows that are at less than 1kb from each other are fused into one CpG island. This procedure leads to a slightly lower number of CGIs compared to UCSC (40,284 vs 48,000). Importantly, a higher proportion of these CpG island are unmethylated in dog sperm (56% for this method vs 46% for the UCSC tracks), therefore reducing the number of false positives. Nevertheless, the two sets of CpG island largely overlap, and using these two different sets of CGIs gave the same results in dogs (fig. S5). I therefore applied this method to call 30,052 CGIs in the genome of the barn owl.

DNA methylation and hypomethylated regions in dog sperm were retrieved from the bi-sulfite sequencing data in Qu et al. (2018).

CGIs were then seperated between those that overlap the promoter of a protein-coding gene (cCGIs) and those that do not (oCGIs). Transcription start sites where taken from the annotation of dog genes from the ucsc (link), and the annotation of barn owl genes from the ncbi (link). Because for a given gene multiple transcripts can be annotated, I only retained one transcript per gene by selecting the longest transcript for which the coding sequence starts with an ATG codon and ends with a stop codon. CGIs were classified as cCGIs if their center was at less than 2kb from an annotated promoter, and classified as oCGIs otherwise. Similarly, a hotspot was found to be in a CGI if at less than 2kb from its center, and outside of CGIs otherwise. Finally, a CGI was classified as hypomethylated if its center was at less than 2kb from a hypomethylated region, and methylated otherwise.

### Repeated element annotations

The repeats in the dog genome were retrieved from the UCSC Table Browser and annotated using Repeat-Masker. I annotated the repeats in the owl genome using EarlGrey, an automated pipeline that identifies *de novo* transposable element families with RepeatModeler and curates consensus sequences with various algorithms (Baril et al., 2024). EarlGrey then annotates these sequences on the genome alongside simple and low-complexity repeats with RepeatMasker (Baril et al., 2024). I used the Archosauria/Testudines database of transposable elements (TEs) from DFAM (Storer et al., 2021) as a reference to identify *de novo* transposable element families and ran EarlGrey with default parameters.

### Computing local increase in CO rate

For each feature, except for repeated elements, I calculated the number of COs that fell within 1.5 kb of its center. The number of CO was then divided by the total number of base pair of the sequences in which they have been called (3kb times the number of features). To control for regional CO rates, I also calculated the number of COs in flanking sequences, which I defined as the sequence between 5 and 30 kilobases (kb) upstream and downstream the feature center. Then, I masked CGIs and hotspots (±1.5*kb*) from these control sequences using *bedtools subtract* (Quinlan and Hall, 2010). Again, I divided this number of COs by the total number of base pair of the control sequences. I then computed the local increase in CO rate as the log_2_ ratio between the CO rate in the features and that of the flanking regions.

For repeated elements, I applied the same procedure as above, except that I kept the whole repeat ±1.5*kb* rather than just the 3kb region around the center. Likewise the control sequences were defined as sequences between 5 and 30 kilobases (kb) away from the closest repeat boundary upstream and downstream of the repeat. Finally instead of removing CGIs and hotpots from the flanking sequences, I removed all other repeats from the same family.

Confidence intervals were obtained by repeating either procedure on 200 bootstrap replicates of the CO events and taking the 5*^th^* and 195*^th^* ordered values of local recombination increase as the confidence interval.

### Matching distributions

At multiple instances, in this study, I compared the local increase in the CO rate of features of interest to a control set of other features. I selected these control features so that either the CpG*_o/e_* ratio distribution or the methylation level matched that of the features of interest. To do so, I approximated the target distribution of CpG*_o/e_*/methylation levels with 40 quantiles. For each quantile, I resampled the features with replacement from the control set until the number of features matched that of the features of interest in the given quantile. The more different the two distributions are, the more a feature will be resampled to fill the quantiles. This leads to greater uncertainty in estimating the local increase in CO rate, as illustrated in fig. S3B.

## Data availability

The position of dog crossovers, CGIs, hotspots, and annotation of the barn owl repeated elements were deposited at https://doi.org/10.5281/zenodo.20747856. All codes necessary to reproduce this study are available at https://gitlab.in2p3.fr/julien.joseph/sexhotuse.

## Acknowledgments

I wish to thank Frédéric Baudat for very helpful discussions and Anaïs Duhamel for help with the figures. I am also grateful to Alexandros Topaloudis and Frédéric Baudat for helpful comments on previous versions of this manuscript. Crossover calling was performed using the computing facilities of the CC LBBE/PRABI.

