## Supplementary material for "Sex differences in PRDM9-independent fine-scale recombination patterns"

### Supplementary material from: Sex differences in PRDM9-independent fine-scale recombination patterns

**Julien Joseph**[1]

[1]Université Lyon 1, CNRS, ENTPE, LEHNA UMR 5023, Villeurbanne, France

June 18, 2026

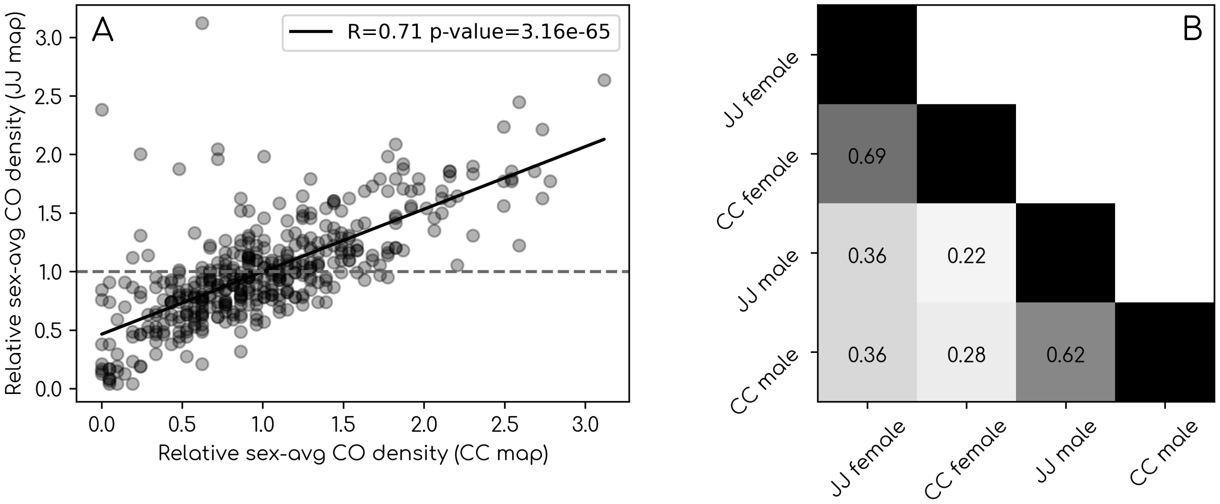


**Figure S1:** The different types of CGIs and their overlap with LD-based hotspots. A) Correlation between sex-averaged recombination rates from this study (JJ) and those from Campbell et al. (2016) Each dot represent a 5Mb window, and the black lines corresponds to a simple linear regression. Dashed horizontal line corresponds to the genome average. B) Correlation matrix representing the pairwise pearson correlation coefficient between sex-specific 5Mb-scale recombination maps from this study and that of Campbell et al. (2016).

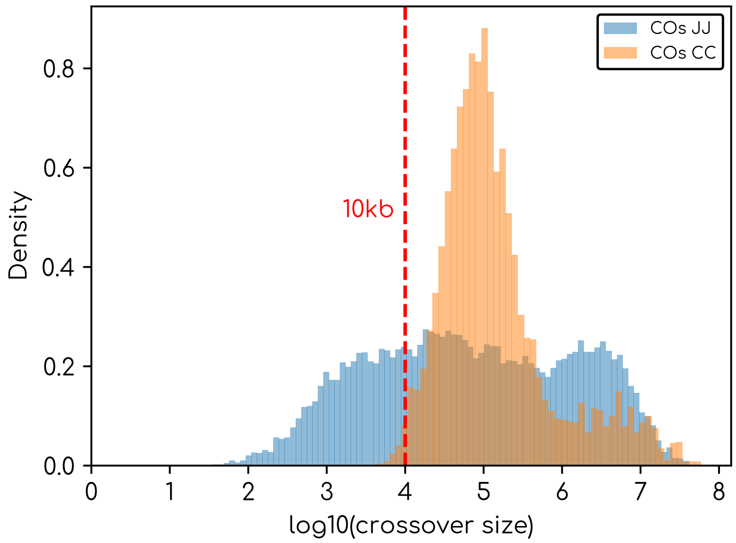


**Figure S2:** Inter-marker distance (or size) distribution of CO events inferred in this study (JJ, blue) and those of Campbell et al. (2016) (CC, orange). Vertical dashed red line indicates COs with a inter-marker distance of 10kb.

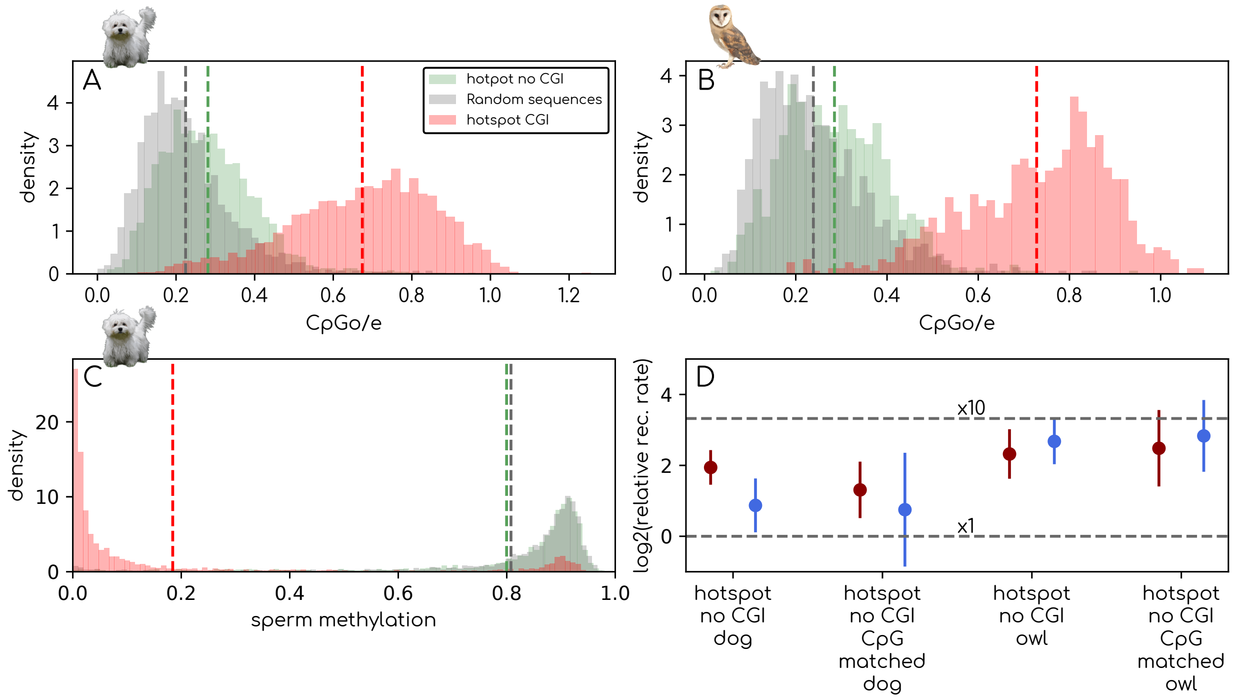


**Figure S3:** A,B) Distribution of $CpG_{o/e}$ ratio in the dog's (A) and owl's (B) non-CGI hotspots. $CpG_{o/e}$ is computed as the number of CpG dinucleotides divided by the product between the number of guanine and cytosine in a 3kb region centered around the CGI. C) distribution of dog sperm methylation level in non-CGI hotspots. Methylation level is the percentage of methylated CpG dinucleotides in in a 3kb region centered around the CGI. Percentage of methylation of individual CpG dinucleotides were obtained from the bisulfite sequencing experiment of Qu et al. (2018). D) $\log_2$ of the relative recombination rate in non-CGI hotspots versus flanking regions in dogs and barn owls. Relative CO rate was computed as the ratio between the number of COs per base pairs in a 3kb region around the feature center and the number of COs per base pairs in the flanking regions defined as above. Error bars correspond to 95% confidence interval obtained by bootstraping CO events (see more details in Material and Methods). CpG matched CGIs are a set of oCGIs selected so that their $CpG_{o/e}$ distribution matches that of cCGIs. Meth. matched CGIs are a set of oCGIs selected so that their sperm methylation level distribution matches that of cCGIs.

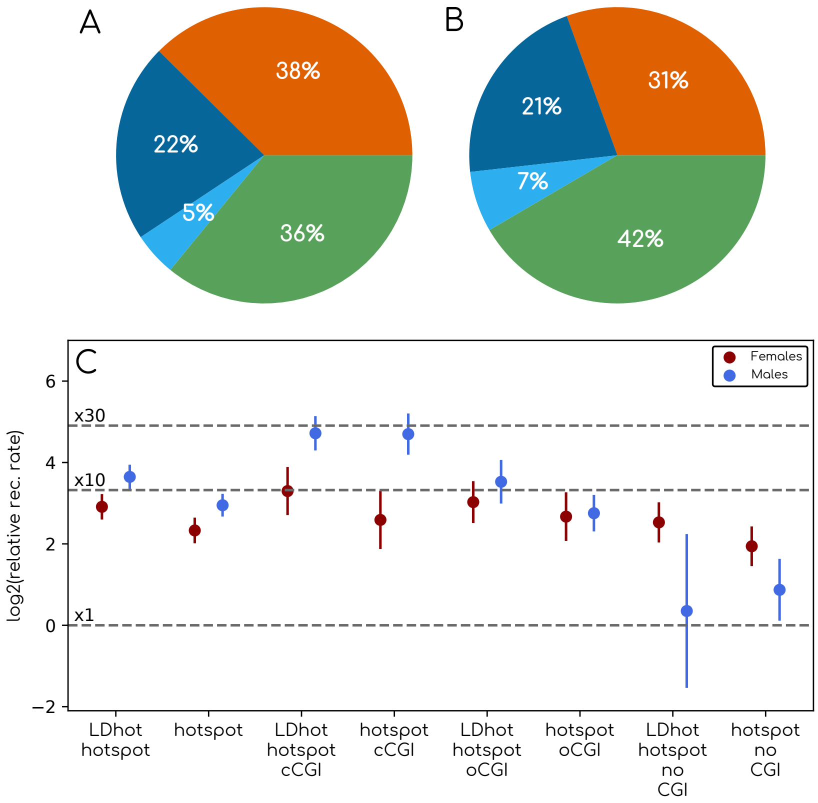


**Figure S4:** A,B) Overlap between LD-based hotspots and the different categories of CGIs in dogs for LD hotspots inferred with LDhot (A) and those used in this study (B). C) Comparison between the $\log_2$ of the relative recombination rate in features versus flanking regions in dogs and barn owls for LD hotspots inferred with LDhot and those used in this study. Relative CO rate was computed as the ratio between the number of COs per base pairs in a 3kb region around the feature center and the number of COs per base pairs in the flanking regions defined as above. Error bars correspond to 95% confidence interval obtained by bootstraping CO events (see more details in Material and Methods).

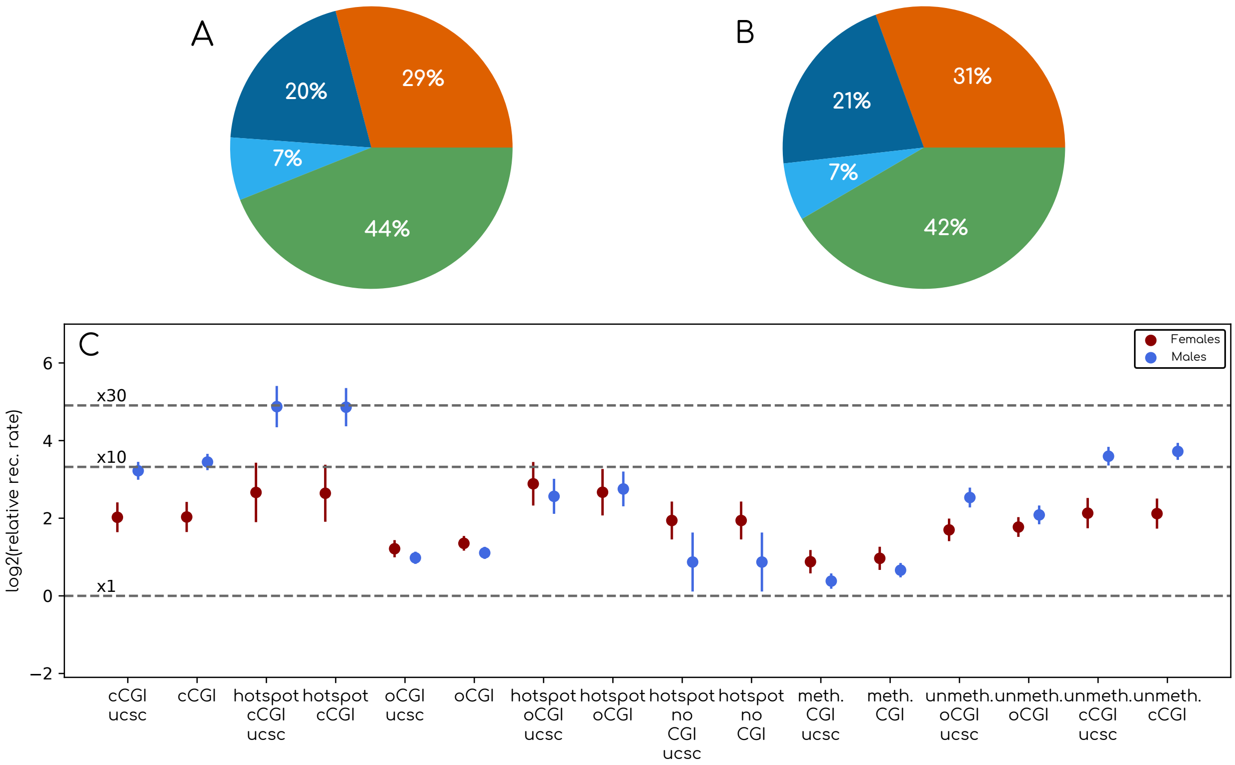


**Figure S5:** A,B) Overlap between LD-based hotspots and the different categories of CGIs in dogs for UCSC CGIs (A) and those used throughout this study (B). C) Comparison between the $\log_2$ of the relative recombination rate in features versus flanking regions in dogs and barn owls for UCSC CGIs (A) and those used throughout this study (B). Relative CO rate was computed as the ratio between the number of COs per base pairs in a 3kb region around the feature center and the number of COs per base pairs in the flanking regions defined as above. Error bars correspond to 95% confidence interval obtained by bootstraping CO events (see more details in Material and Methods).
